# *fea*miR: Feature selection based on Genetic Algorithms for predicting miRNA-mRNA interactions

**DOI:** 10.1101/2020.12.23.424130

**Authors:** Eleanor C. Williams, Anisoara Calinescu, Irina Mohorianu

## Abstract

microRNAs play a key role in RNA interference, the sequence-driven targeting of mRNAs that regulates their translation to proteins, through translation inhibition or the degradation of the mRNA. Around ~ 30% of animal genes may be tuned by microRNAs. The prediction of miRNA/mRNA interactions is hindered by the short length of the interaction (seed) region (~7- 8nt). We collate several large datasets overviewing validated interactions and propose ***fea*miR**, a novel pipeline comprising optimised classification approaches (Decision Trees/Random Forests and an efficient feature selection based on embryonic Genetic Algorithms used in conjunction with Support Vector Machines) aimed at identifying discriminative nucleotide features, on the seed, compensatory and flanking regions, that increase the prediction accuracy for interactions. Common and specific combinations of features illustrate differences between reference organisms, validation techniques or tissue/cell localisation. *fea*miR revealed new key positions that drive the miRNA/mRNA interactions, leading to novel questions on the mode-of-action of miRNAs.

## 1 Introduction

Recent improvements to sequencing technologies enabled a wide range of biological questions, through larger, more diverse experiments [1]. One area that thrived is the prediction of regulatory interactions [2]; further advances, e.g. improving prediction accuracy, could lead to a more precise description of biological processes [3]. A key regulatory role in gene networks is played by non-coding RNAs; a class of small noncoding RNAs, microRNAs (miRNAs), ~ 20 —22 nucleotides long, influences, in animals and plants, the protein output through transcriptional and post-transcription mRNA regulation [3, 4]. Although ~ 30% of genes in mammalian cells could be regulated by miRNAs [5], the accuracy of target-prediction is still low [6]. However, the increasing number and diversity of validated miRNA/mRNA interactions, coupled with information on the localisation of miRNAs and mRNAs, has the potential to reveal novel discriminative features, that may lead to an increase in the accuracy of predictions.

Most prediction methods of regulatory interactions summarise data-mining results into rule-based models [7, 8, 9, 10]. This approach carries the risk of overfitting models to specific sets of observations, while minimising their robustness and portability to new data across organisms [11]. Current tools for predicting miRNA/mRNA regulatory interactions are based on string-matching approaches [12, 13]. Accepted criteria focus on perfect complementarity of the miRNA seed region (~7-8nt from the 5’ end of the mature miRNA) to its target mRNA. MiRanda [14] developed on *D. melanogaster* filters pairs according to seed matches, free energy and conservation across the *Drosophilidae* clade. miRanda-mirSVR [15] is an extension based on support vector regression-generated scores for quantifying the strength of the interaction using site accessibility, position of target, and flanking content as predictive features. TargetScan [16] requires some degree of complementarity on the compensatory region (centred sites) in addition to the perfect complementarity on the seed region, to limit the number of false-positives; additional conservation-based filters are also used. DIANA-microT-CDS [17] outputs predicted target location and binding type, and introduces conservation for either the miRNA, mRNA or the interaction, by identifying important features from PAR-CLIP data.

These tools achieve limited accuracy and sensitivity (e.g. *se*= 0.643, TargetScan); the restricted set of features considered in rule-based approaches contribute to a large number of false-positives [18] and requires regular updates to incorporate the substantial increase in miRNA annotations [19] and validated interactions [20].

To improve the accuracy, sensitivity and specificity for predictions of miRNA/mRNA interactions, we focus on true targets as a starting point, and characterise the positional-nucleotide content of (i) the miRNA seed region, (ii) the miRNA compensatory site, and (iii) the upstream and downstream mRNA flanking regions, to discriminate between validated and non-validated miRNA/mRNA interactions. We investigate the distributions of discriminative features across model organisms, validation approaches, and tissue localisation. We assess statistical approaches and classical ML models (Decision Trees, Random Forests and SVMs). SVMs are enhanced with novel feature selection approaches (using embryonic Genetic Algorithms, for a fast and optimal search of the feature space), leading to a higher accuracy than existing approaches.

## 2 Results

### 2.1 Statistical results on positional features

In line with existing approaches [16, 17], we focused on single and di-nucleotide positional features processed in a one hot encoding (SM1). We assessed, using the χ^2^ test on all features per position and the Fisher exact test to evaluate individual categories, the differences between the positive and negative set, validated and non-validated interactions, respectively [20]. We observe several discriminative features concentrated in the seed and compensatory regions (Figure 1a, *H. sapiens*); however, using all interactions, none were observed on the flanking regions.

**Figure 1.**
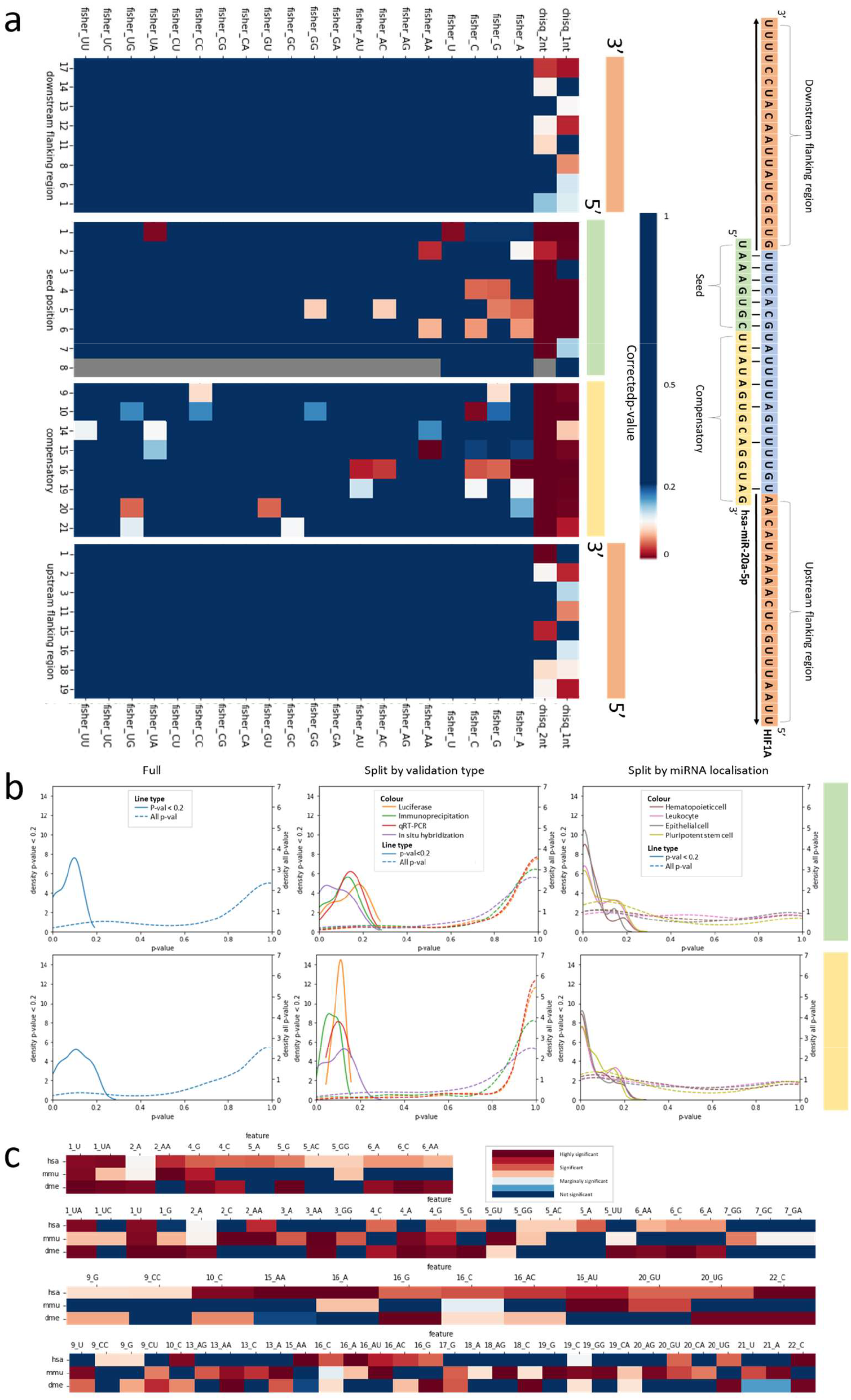
Overview of statistical significance of single and di-nucleotide positional features for *H. sapiens, M. musculus* and *D. melanogaster*. [a] Heatmaps illustrating χ^2^ and Fisher exact BH adjusted p-values, assessed on the frequency distributions of single and di-nucleotide positional features, testing differences between validated and non-validated interactions. From left to right, we present downstream flanking, miRNA seed, miRNA compensatory and upstream flanking regions. When all interactions are considered, significant features are concentrated on the seed and compensatory regions e.g. position1_U (enriched U at position 1, seed region), position10_C and position16_A. [b] Density plots on distributions of p-values for Fisher exact tests, on full positive and negative sets, for seed (top) and compensatory (bottom) position-nucleotide combinations, with Benjamini-Hochberg corrections; the LHS and RHS y-axes illustrate the scale for the raw and BH adjusted p-values, respectively; the line type indicates all features (dotted line) or features with p-value<0.2, significant or marginally significant (solid line). The positive set is also split by validation type and localisation. The distributions of p-values for the first two categories are similar; however we observe a significant shift towards near-zero p-values if cell type is considered. [c] Heatmaps illustrating the conservation of significant features between *H. sapiens*, *M. musculus* and *D. melanogaster*. The features shown are, from top to bottom, significant seed features in *H. sapiens*, significant seed features in *H. sapiens* or *M. musculus*, significant compensatory features in *H. sapiens* and significant compensatory features in *H. sapiens* or *M. musculus*. We see the majority of features are conserved across 1 or more species.

**Figure 2.**
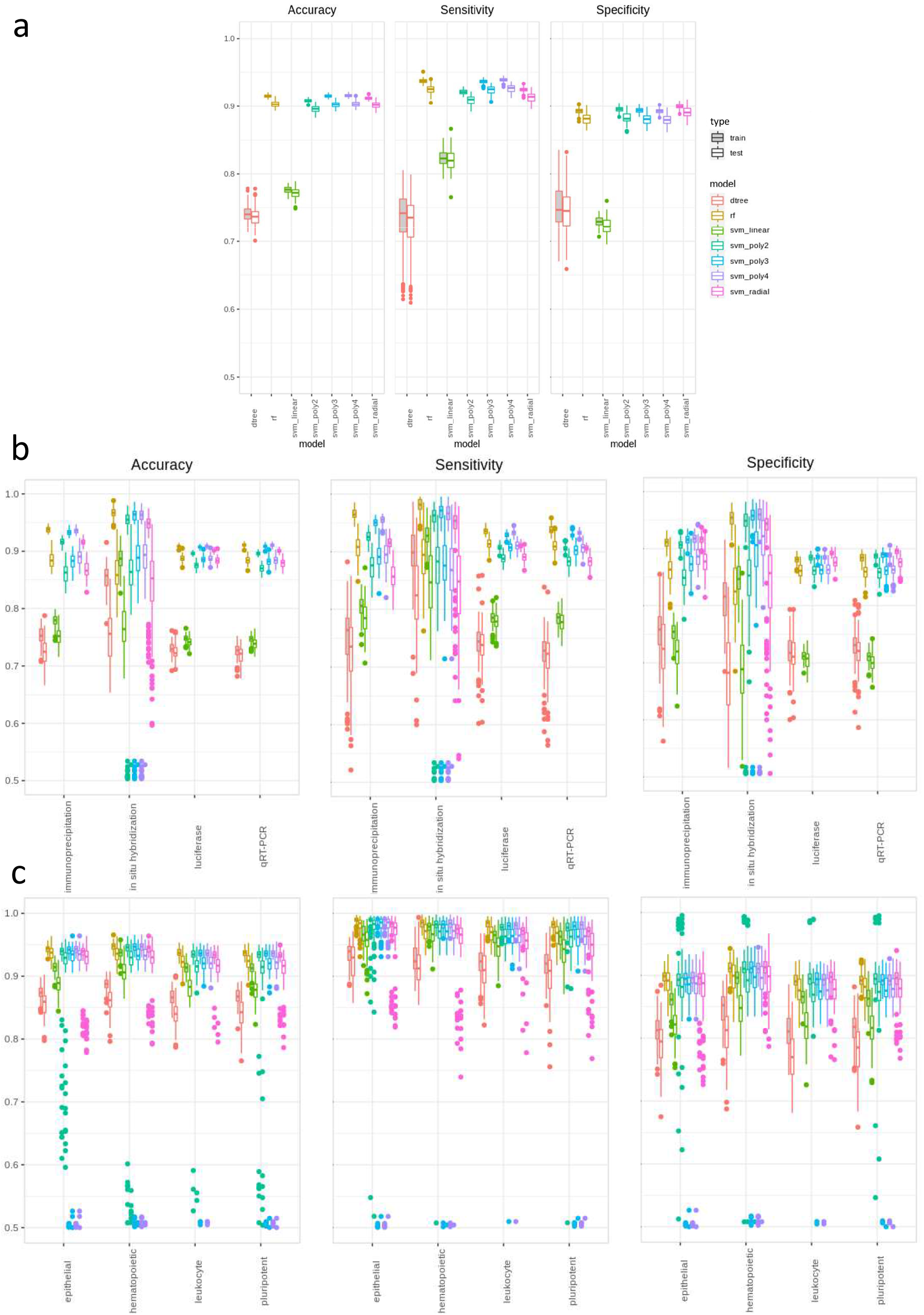
Overview of performance of classifiers using seed features on *H. Sapiens*. Performance of Decision Trees, Random Forests (ntree=30) and SVMs (with several kernels) across 100 subsamples of the full positive and negative set. Each subplot shows the distribution of test and training accuracies, sensitivities and specificities for [a] all interactions, [b] interactions split per validation type and [c] interactions split per localisation. For all interactions, the DTs and linear kernel SVMs show consistently lower performance, while the RFs and SVMs with poly4 kernels achieve the best results. A higher variability is observed for the validation types, corroborated with larger differences between the training and testing measures, for the in-situ hybridization, which suggest an overfitting due to insufficient information. The sensitivity results for the localisation split are remarkably tight and high, supporting the hypothesis that different subsets of features may be functionally-relevant across cell types.

To focus the analysis, we employed additional information from miRTarBase related to predictionconfidence (the Functional MTI (miRNA–target interactions) and weak Functional MTI categories) and validation approaches. Using the former, no single or di-nucleotide features had p-value<0.1. We observe higher statistical significance for strong-validation features (out of 502,652 interactions for *H. sapiens,* 10,754 were labelled as non-weak Functional MTI). On this restricted set a higher number of features are statistically significant (χ^2^ tests), on the seed, compensatory and flanking regions. On the former we observe significant adjusted *p*-values for Fisher exact tests on specific position-nucleotide combinations e.g. position1_U, pair2_AA, position10_C, pair15_AA and position16_A.

Conserved features on *H. sapiens, M. musculus* and *D. melanogaster* make up 36% of significant features e.g. position1_U, pair1_UA, position2_A and position4_G. Based on pairwise comparisons 46% of seed features are conserved between *H. sapiens* and *M. musculus,* e.g. pair5_GG and position2_AA, and 69% conserved between *H. sapiens* and *D. melanogaster,* e.g. position4_C, position6_C and position5_G. Species-specific features are observed outside the seed region, e.g. pair15_AA for *H. sapiens,* pair16_AU for *M. musculus* and pair20_UG for *D. melanogaster.*

Additional information retrieved from miRTarBase [20] relates to validation; to assess the differences in discriminative features introduced by this covariate, we focused on subsets with ≥ 40,000 entries including luciferase reporter assay, immunoprecipitation, qRT-PCR, and in-situ hybridization. We identified different feature-sets depending on this covariate (Figure 1b); several features were identified as discriminative for all experiment types across interaction regions e.g. position1_U and pair1_UA on the seed and position22_C and pair16_AU on the compensatory region. However some features were specific to validation types, e.g. pair1_UA is significant in luciferase and immunoprecipitation but only marginally significant for qRT-PCR. We also observed 8 features specific only to in-situ hybridization, e.g. position4_G, position9_G and pair20_UG (Supplementary figure 1). These results underline differences in the types of interactions (and selected features) captured by the validation experiments.

To further assess the variation in the selected feature-set we introduced an additional covariate, the localisation of miRNA across several cell types [21]; we focused on cell types with sufficient validated entries i.e. hematopoietic cells, epithelial cells, leukocytes and pluripotent stem cells (104, 129, 138 and 141 miRNAs respectively). Figure 1b illustrates significant differences in p-value distributions on the full set, the set split by validation and the entries split by cell type. This suggests the existence of a functional signature for validated miRNA/mRNA interactions; on the seed region 5.4% of discriminative features were shared across all 4 localisation categories, 20.3% by 3, 28.3% by 2 and 32.4% selected by only 1 localisation category; on the compensatory region, we see 7.O% of features are shared 4-way, 21.1% by 3, 27.2% by 2 and 44.7% by 1. A noteworthy difference is observed for the flanking regions, no features are selected 4-way and 69.4% and 58.4% of discriminative features selected only by one localisation category for the upstream and downstream flanking regions respectively. Based on these results, we infer that miRNA/mRNA interactions may be functionally cell type-specific.

### 2.2 Classical classifiers and the selection of discriminative features

To characterize interactions between features and determine optimal combinations for discriminating the positive and negative set, we applied several classical classification approaches: Decision Trees (DTs), Random Forests (RFs), Support Vector Machines (SVMs); each applied with default parameters, and subsequently used with selected, discriminative features.

Individual DTs, trained on 100 different subsamples with equal numbers of validated and nonvalidated interactions on *H. sapiens*, using only seed features, achieved a mean training accuracy of 0.741, and test accuracy of 0.737. While the the two values are comparable, this performance is low and suggests underfitting; the features selected on top levels include position6_A and pair1_UA (also identified as statistically significant), pair3_GG, pair6_GU, pair1_UA and pair3_CA.

To overcome the limitations of DTs we tested RFs; the number-of-trees hyperparameter was tuned using cross-validation *(ntree*= 30). The mean accuracies were 0.915 and 0.904 on training and test, respectively. In contrast to DT, the sensitivity also increased; mean training and test values observed were 0.937 and 0.924 respectively (from 0.724 and 0.721 for DT). The most discriminative features included pair1_UA, pair3_GG (shared with DT), position1_U, pair5_GG and position5_A.

To further enhance the classification, we used SVMs with tuned kernels (polynomial kernels and a radial basis one). The polynomial degree 4 kernel was chosen based on its consistent high performance (mean training and test accuracies 0.915 and 0.902 across 100 runs on different subsamples and mean sensitivities 0.936 and 0.923).

For all validation types, except in-situ hybridization, RFs and SVM outperform DTs; the observed accuracy ranges are [0.85, 0.9] and sensitivity [0.9,0.95]. The performance for in-situ hybridization highly varied, suggesting overfitting. Across cell types, RFs and SVM outperform DTs. The RFs perform better when training examples are split by validation type. These results support the hypothesis of functionally-specific features.

### 2.3 Targeted feature selection using (embryonic) Genetic Algorithms

Further to using the well-established classifiers, on all features, for direct prediction of miRNA/mRNA interactions, we exploited feature selection approaches on DTs, RFs and SVMs.

Using a DT voting scheme on features (Supplementary method 1) for the top 10 levels of 100 DTs trained on the full positive set and different subsamples of the negative set, we observe 5 features consistently selected (87-99 times/100 runs): position1_U, pair1_UA, position6_A, pair3_GG and pair6_GU. The first 3 features were statistically significant; however pair3_GG and pair6_GU were not.

Using the in-built Gini index node impurity measure from RFs, the top 5 discriminative features are position1_U, pair1_UA, pair3_GG, pair5_GG and position5_A. 4 features overlap with the DT ranking (pair5_GG was the 6th most commonly selected feature); position5_A was not chosen by DT, however it was statistically significant.

Due to time complexity, Forward Feature Selection was only performed once and the feature ranking was cut off at 20. For FFS position1_U was selected as the top feature, followed by pair3_GG and pair5_GG (RF), pair7_GG (selected in the top 20 by DT and RF) and pair4_UC not identified as significant yet. In Figure 3d, we show that adding up to 20 features increases test accuracy and sensitivity, however we observe a trade-off on sensitivity/accuracy and specificity if more than 15 features are considered.

**Figure 3.**
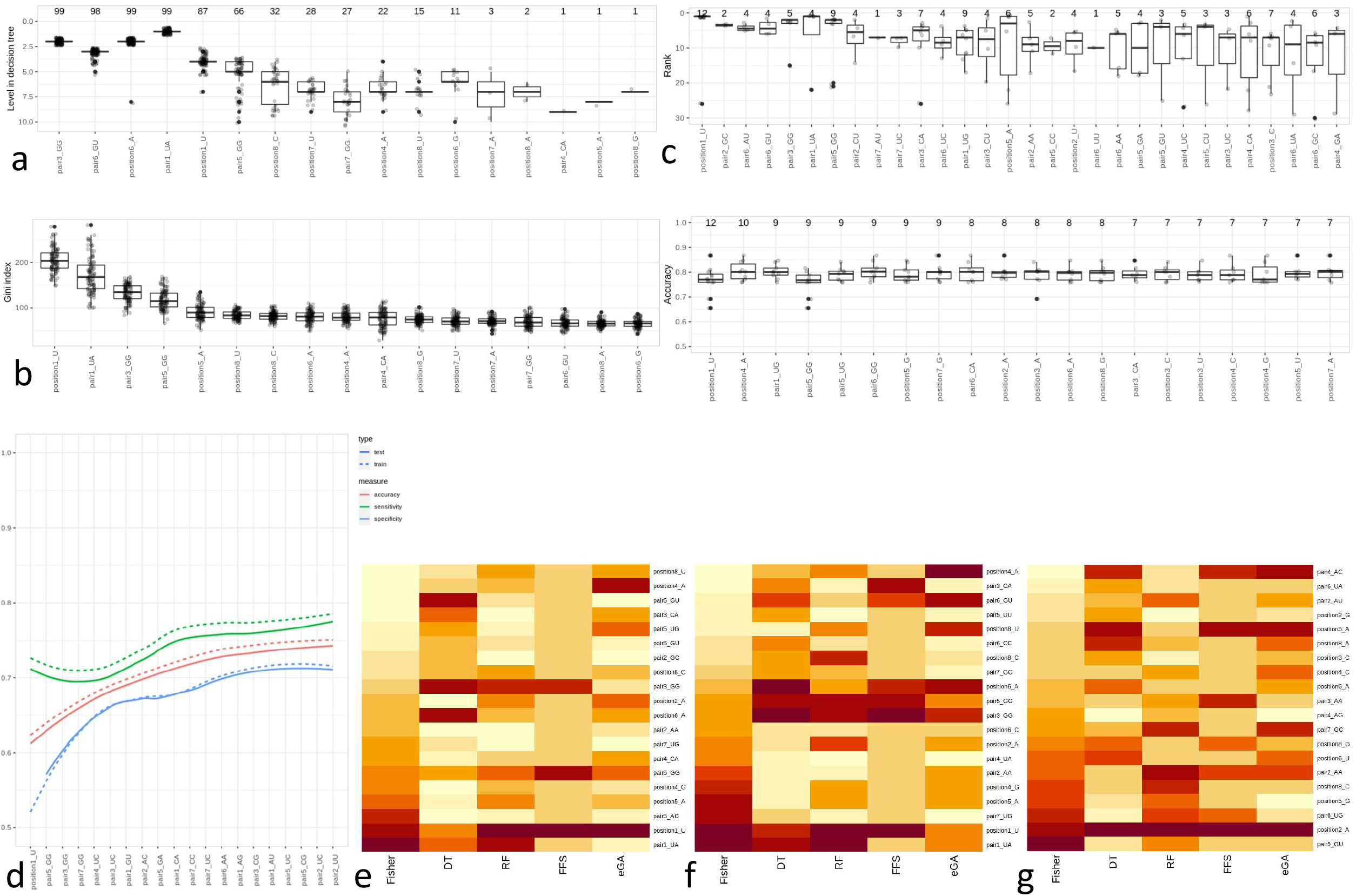
Summary of feature selection approaches on *H. sapiens*. [a] Boxplot showing distribution, on DTs, over 100 subsamples, of depth from the root for each feature (considering only features appearing in the top 10 levels); we observe a sharp and sudden decrease for the feature frequencies, shown on top of every boxplot, combined with a high variability in rank. [b] Boxplot showing distribution of mean decrease in Gini index of features across 100 runs on different subsamples, ordered by cumulative mean decrease in Gini. The voting scheme in the RFs control and minimise the variability in information content across features. [c] Summary of features selected by eGA on full positive and negative sets. Distribution of ranks (top) and accuracies (bottom) across 100 subsamples for the training data). On top of the boxplots we present the number of selections per feature. [d] Improvement in performance when adding features, ordered on the x-axis, selected by FFS. The line type indicates the training (dotted line) and test (continuous line) accuracy, sensitivity and specificity for the first 20 features. [e] Heatmap showing the discriminative power of features across the 5 feature-selection methods on the full positive set: Fisher exact p-values, DT voting scheme, RF cumulative mean decrease in Gini, FFS and eGA. The top 10 features across methods were included; all scores were quantile normalised on all 144 features per method, for comparability. The stronger features are shown in darker red. [f,g] Discriminative power heatmaps for positive examples validated by luciferase experiments [f] and on miRNAs localised to pluripotent stem cells [g].

We also tested the novel eGA approach [22] as it further explores the links/connections between features; features identified include pair6_GU and pair2_GC which acquire highest ranks when chosen but are not preferred often. The features selected as most discriminative are position1_U, pair5_GG, pair1_UG, position4_A and position2_A. These and several other high-rank features are consistent throughout multiple prediction methods.

A summary of the features selected across methods is shown in Figure 3e; we observe that each feature selected by eGA is also significant for some other method, unlike the other approaches which choose also specific features e.g. pair5_AC identified using Fisher exact tests or pair6_GU in DTs. The novel eGA captures a consistent set of highly ranked features, e.g. pair1_UA, position1_U and position5_A and pair1_UG. SVMs trained on the features selected by the eGA achieve stable accuracy around 0.8, higher than on the features chosen by FFS.

Concentrating on the identification of an optimal set of features, the output of *fea*miR for the *H. sapiens* dataset, with all interactions considered, is an SVM classifier on polynomial degree 4 kernel; the selected, consistent features include position1_U, position6_A, pair1_UA, pair5_GG, position2_A and pair3_GG. The generic output of feamiR is an optimal classifier for discerning validated vs unvalidated interactions, and a ranked selection of discriminative features for this task.

To further assess the interactions between features we partitioned the interactions on the validation technique. Figure 3f summarises the selected features for luciferase validation (similar results on other validations types are illustrated in Supplementary figure 3). Based on statistical tests, position1_U and pair1_UA are consistently identified across validations and feature selection approaches, alongside position6_A, position5_A, pair5_GG and pair3_GG also selected for the full positive set. Specific features stable on subsets of validations include position2_A, discriminative for luciferase, immunoprecipitation and qRT-PCR, but not for in-situ hybridization, or position4_G discriminative only for in-situ hybridization. The presence of both consistent and specific features suggests that combining feature selection approaches, we can increase the robustness of the selection and extend the set of discriminative features.

More variability in the subset of selected features is observed across localisation categoriese.g. in pluripotent stem cells we position2_A is robustly identified by all approaches but not chosen in any other cell types; other examples include.position7_G in hematopoietic cells, position8_A in leukocytes and position1_A in epithelial cells. Discriminative features across the whole positive set, e.g. position1_U are present across cell types.

### 2.4 *fea*miR: an R Package for the selection of discriminative features, predictive of miRNA/mRNA interactions

*fea*miR integrates all options described in previous sections, for determining the discriminative features, predictive of miRNA/mRNA interactions; it is available as a stand-alone CRAN package and comprises a pipeline (detailed in Figure 4) taking as input a set of miRNAs, a set of mR-NAs (3’ UTRs) and an interactions database, partitioned into validated pairs (‘positive’ set) and non-validated pairs (‘negative’ set); the latter consist of (miRNA, mRNA) tuples, with seed or compensatory complementarity which are not present in the validated set. For both sets, all single and di-nucleotide features are considered for [a] statistical assessment and [b] classification and feature selection.

**Figure 4.**
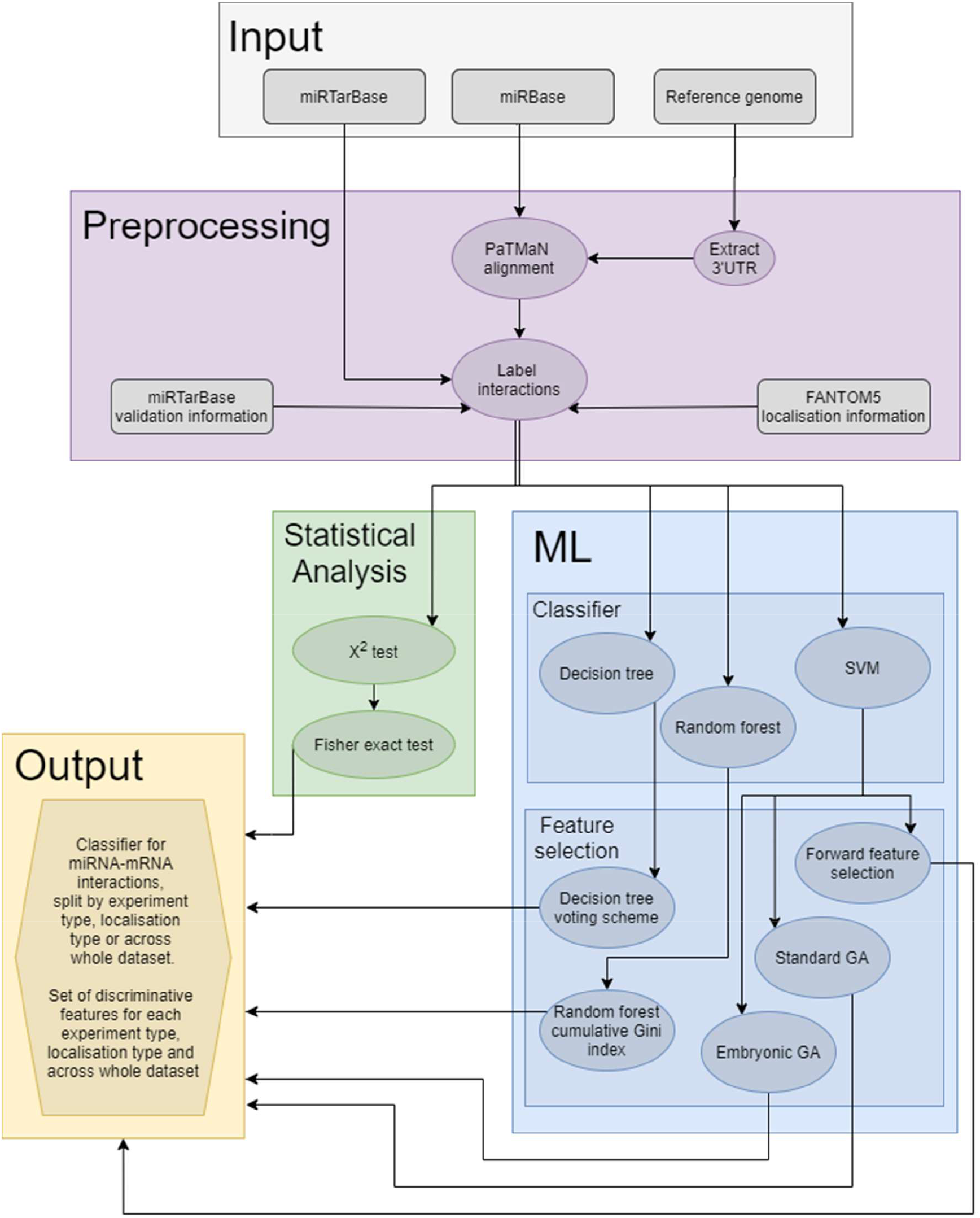
*fea*miR workflow diagram. The processing pipeline handles miRNA and 3’UTR sequences. These, in conjunction with external information (e.g. miRTarBase or FANTOM5), produce a series of discriminative features, ordered on their predictive power, and a classifier with high predictive power. It comprises a preprocessing section and two data analysis sections, based on statistical and Machine Learning approaches; the latter are augmented using feature selection approaches based on Genetic Algorithms.

The preprocessing steps i.e. the formatting of the 3’ UTRs and generating the miRNA/mRNA pairs, are performed in Python and integrated through external system calls. Preliminary statistical analyses are then performed (χ^2^ and Fisher exact tests as detailed in Results). Additional, external information, such as validation type or localisation of the miRNA/mRNA pair can be used to further focus on discriminative features (the analysis proceeds on individual subclasses only where ≥ 40 (or another user-input threshold) entries for a single validation type are available).

Several classifiers (DTs, RFs, SVMs) coupled with feature selection approaches, (based on variable importance, Forward Feature Selection or GA/eGA) are also included in *fea*miR; with a higher accuracy and sensitivity, these identify positional nucleotide features which are highly discriminative for separating the positive and negative entries. Additional functions for fine-tuning classifierspecific hyperparameters such as the RF number-of-trees *(selectrfnumtrees*) and SVM kernels *(se-lectsvmkernel*) are available; the *runallmodels* function uses these hyperparameters to generate a summary of how each model performs across all 100 subsamples. Optimal values are inferred through cross-validation on the input data; the users can set (some of) these parameters based on external knowledge. Further guidelines and examples for using this package are presented on github: https://github.com/Core-Bioinformatics/feamiR.

## 3 Discussion

### 3.1 Accuracy, Sensitivity and Specificity of *fea*miR versus existing approaches

Either on the whole dataset or split by validation or localisation, we observe on 100 iterations of k-fold cross-validations that SVMs and RFs are optimal models. With the exception of in-situ hybridization, SVMs and RFs perform similarly; because of the risk of RFs identifying local optima, for the biological interpretation, we prefer SVMs. For in-situ hybridization and on several localisation categories, RFs outperform SVMs due to a few outlier runs from SVMs; excluding these leads to higher SVM-accuracies. The performance of RFs on specific localisation categories is higher than across the combined set, e.g. for epithelial cells *μ_acc_*= 0.94 (σ= 0.0097) compared to *μ_acc_*= 0.91 (σ= 0.0066) on the whole dataset (the accuracies and sensitivities across classes are summarised in Table 1).

**Table 1.** Summary of accuracy, sensitivity and specificity achieved using *fea*miR across different cell types (epithelial cell (EC), leukocyte (Le), pluripotent stem cell (PSC), hematopoietic cell (HC)) and validation techniques (immunoprecipitation (Im), in-situ hybridization (ISH), luciferase (Lu), qRT-PCR (qRT). Across all case studies, the classifier using enhanced, selected features outperforms existing approaches, with minimal variation for the quality of prediction. In particular, the highest accuracy and sensitivity combination is achieved using RFs on miRNAs localised to epithelial cells, compared to the lower performance for in-situ hybridization. The tightest range is observed for SVMs on interactions validated using luciferase and qRT-PCR experiments.

| Dataset | All SVM | Localisation |  |  |  | Validation |  |  |  |
| --- | --- | --- | --- | --- | --- | --- | --- | --- | --- |
|  |  | EC RF | Le RF | PSC RF | HC RF | Im SVM | ISH RF | Lu SVM | qRT SVM |
| Acc range | 0.89-0.92 | 0.91-0.97 | 0.89-0.95 | 0.89-0.99 | 0.90-0.97 | 0.88-0.92 | 0.79-0.93 | 0.86-0.90 | 0.86-0.89 |
| Acc $\mu$ | 0.91 | 0.94 | 0.92 | 0.93 | 0.94 | 0.89 | 0.86 | 0.88 | 0.88 |
| Acc $\sigma$ | <0.01 | <0.01 | 0.01 | 0.02 | 0.02 | 0.01 | 0.02 | <0.01 | <0.01 |
| Se range | 0.91-0.94 | 0.95-1.00 | 0.88-0.99 | 0.89-0.98 | 0.80-0.99 | 0.86-0.92 | 0.77-1.00 | 0.87-0.92 | 0.85-0.91 |
| Se $\mu$ | 0.93 | 0.98 | 0.95 | 0.93 | 0.90 | 0.90 | 0.89 | 0.89 | 0.88 |
| Se $\sigma$ | <0.01 | 0.07 | 0.1 | 0.08 | 0.11 | 0.02 | 0.05 | 0.01 | 0.01 |
| Sp range | 0.86-0.90 | 0.84-0.94 | 0.84-0.94 | 0.82-0.97 | 0.69-0.97 | 0.84-0.93 | 0.72-0.96 | 0.85-0.90 | 0.85-0.91 |
| Sp $\mu$ | 0.89 | 0.90 | 0.88 | 0.91 | 0.84 | 0.88 | 0.95 | 0.88 | 0.87 |
| Sp $\sigma$ | <0.01 | 0.02 | 0.02 | 0.03 | 0.21 | 0.02 | 0.05 | 0.01 | 0.01 |

The classical ML models consistently attain higher sensitivity (and accuracy), both across the whole dataset and split by validation or localisation, than existing models, e.g. miRAW, a deep learning approach [23] reported se = 0.72 and *acc*= 0.78, miRanda [14] reported se = 0.672, and TargetScan [16] registers se = 0.643. RFs and SVMs on all position-nucleotide features on the miRNA seed region markedly improves upon the accuracy and sensitivity of existing techniques (Table 1).

An informative result from feamiR is the priority-ordered subset of position-nucleotide features, particularly effective in differentiating between validated and non-validated interactions e.g. position1_U, position6_A, pair1_UA, pair5_GG, position2_A and pair3_GG are consistently chosen by different feature selection approaches. Training models on a reduced, focused set of discriminative features achieves *acc*= 0.83 and se = 0.86. The selection of features can also be further explored from a biological angle, to understand factors driving the miRNA/mRNA interactions.

### 3.2 Impact of available information on the accuracy of prediction

For *H. sapiens* and *M. musculus,* the miRTarBase dataset comprises a high number of validated interactions (502,652 and 49,830, respectively), leading to sufficiently large positive sets. The interaction dataset for *D. melanogaster* is orders of magnitude smaller (192 interactions) resulting in over-amplified differences between positive and negative sets and overfitting to the positive examples. In addition, this leads to a minimal representative power of the subsamples used for the negative set. For the latter, consequently, the statistical analysis yields a large number of significantly discriminative features between the positive and negative sets *(M. musculus* and *D. melanogaster* in Supplementary figure 1). Subsequently we observe high accuracies *(μ*= 0.91, σ= 0.02) and sensitivities *(μ*= 0.97, σ= 0.02) for RFs and SVMs (Supplementary figure 2) that are likely to indicate overfitting. This underlines the limitations of this analysis on species (subsets) with few validated interactions i.e. the results may be biased by insufficient information and some features may be artificially classified as discriminative.

Unlike many current approaches focused on fixed rules for miRNA/miRNA classification, *fea*miR adapts the feature-set in a information-driven manner; an example of enhanced accuracy and sensitivity is observed when integrating localisation information from FANTOM5 [21]. The differences between cell types induce diverse feature subsets; illustrated in Supplementary figure 4 are the p-values for *χ^2^* tests between nucleotide composition in each pair of localisation categories (averaged over 8 seed positions). Interestingly, we notice biologically related subgroups of cell-types with consistent interaction properties e.g. iPS, embryonic stem cells and pluripotent stem cells, or ectodermal and neuroectodermal cells. However differences between unrelated cell types are significant and support a customised, data-driven set of features which may lead to a more accurate characterisation of localised, functionally relevant, mode of action. The extent of differences observed for *M. musculus*compared to *H. sapiens* cell types underlines the limitations of current datasets and the benefits from more entries and manual curation that could be incorporated into predictions. Examples of localised changes in mode of action for miRNAs have already been described in plants [24].

Interestingly, we observe significant differences in position-nucleotide composition on the flanking regions between cell types. To evaluate the hypothesis on downstream and upstream flanking regions, we examined 50nt regions. In Figure 5a we summarise the number of significant positions per 10nt window. We observe a decrease in the number and significance level of features as we move away from the miRNA complementary region, with a clear drop-off after 40nt in all 4 tested cell types. We also see a decrease in the number of highly significant features after 20nt in epithelial cells and pluripotent stem cells. In line with this remark, at the single cell level we focus on features across 20nt flanking regions.

**Figure 5.**
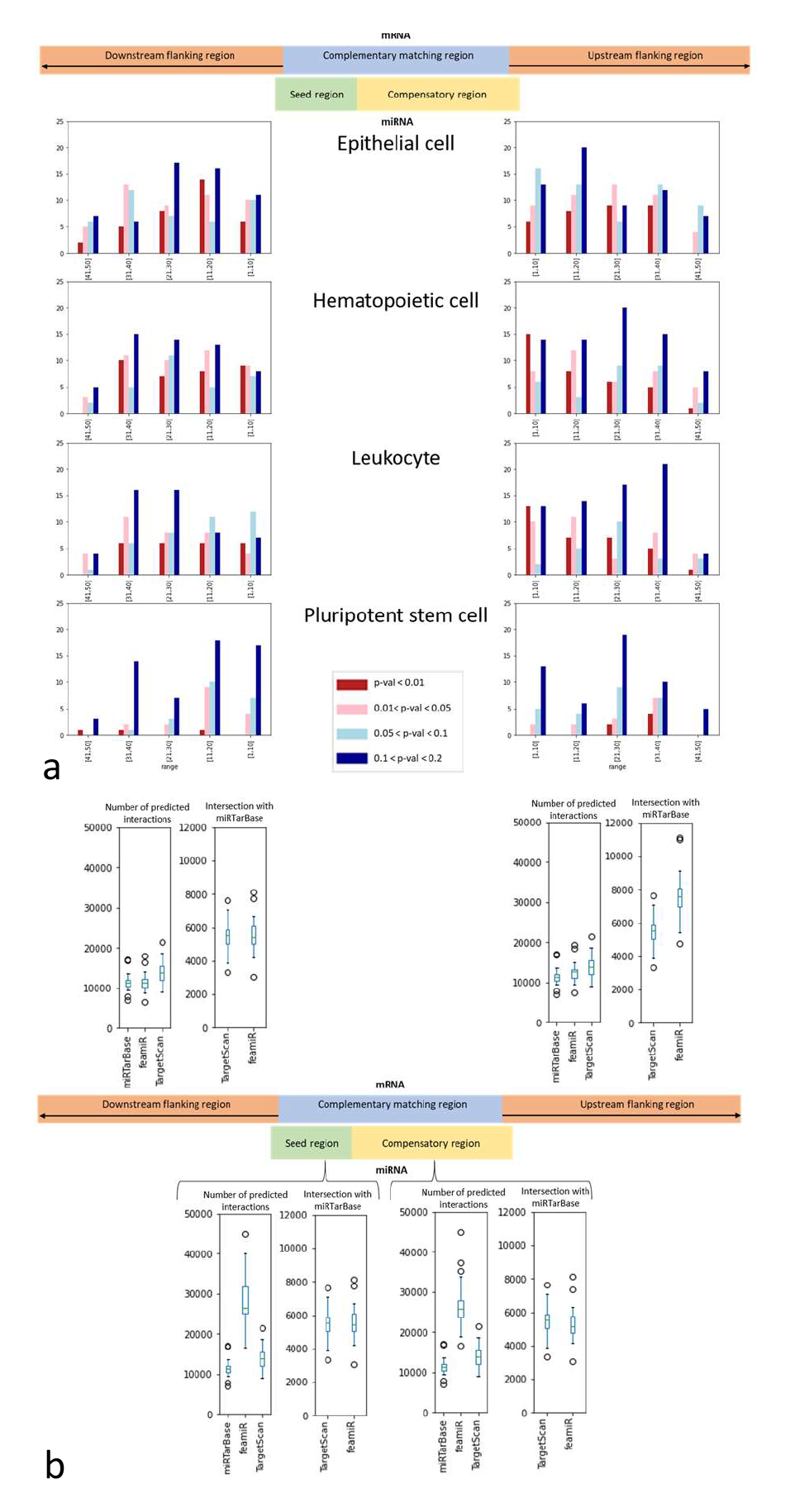
Overview of the discriminative power of flanking features for interactions localised to specific cell types and at single cells level, in *H. sapiens*. [a] Histograms showing the number of significant features and significance levels of features grouped in 10nt bins for the 50nt upstream and downstream flanking regions; these are assessed using Fisher exact tests on epithelial cells, hematopoietic cells, leukocytes and pluripotent stem cells. We observe a decrease in the number and significance levels of features away from the miRNA complementary region, with a clear drop-off after 40nt in all cell types. In addition we notice a decrease in the number of highly significant features after 20nt for epithelial cells and pluripotent stem cells. [b] The distribution of number of interactions predicted by TargetScan, miRTarBase and feamiR (using seed, compensatory, upstream and downstream flanking features) across 19 cells, in a single cell co-expression experiment. We show the number of interactions per prediction tool and the size of the intersection of TargetScan and feamiR with miRTarBase (considered the true positive set). Using seed or compensatory features, *fea*miR predicts more interactions than TargetScan and miRTarBase, however using upstream or downstream flanking features, *fea*miR predicts fewer interactions than TargetScan yet the intersection with miRTarBase is larger, supporting the observed increased sensitivity.

### 3.3 Single cell analysis

To expand on the differences of the miRNA/mRNA mode of action across cell types, we focused on single-cell miRNA-mRNA co-sequencing [25]. Using the sRNA and mRNA quantification from 19 cells, we compared the number of interactions per cell predicted by miRTarBase, TargetScan and *fea*miR, on co-localised miRNAs and mRNAs. Concentrating on seed features, *fea*miR predicts more interactions *(μ_feamiR_Seed_*= 28156, *σ*= 6602) then miRTarBase *(μ_miRTarBase_* = 11471, *σ*= 2444) and TargetScan *(μ_TargetScan_*= 14082, σ= 2858), with a large overlap with TargetScan output (59%), underlining its emphasis on seed complementarity. To balance the high probability of finding near-perfect complementary between the seed-sequence and 3’UTRs for a restricted number of cellspecific miRNAs we analysed also the extended feature set. The conclusion holds for compensatory features; *fea*miR predicts a mean number of interactions *μ_feamiR_comp_*= 26867 (σ= 6680) with 57% intersection with TargetScan.

When flanking features are used, we predict fewer interactions, thus reducing the false-negative rate lower than TargetScan, and also produces a higher true positive rate (shown by the number of miRNA-mRNA pairs in the intersection between miRTarBase and *fea*miR predictions). While there are fewer significant flanking features in the full positive set, we observe more significant differences when localisation is considered, both on cell type and single cell levels. The distribution of number of interactions predicted by TargetScan, miRTarBase and *fea*miR (using seed, compensatory, upstream and downstream flanking features) across 19 cells is shown in Figure 5b.

This suggests that current tools focusing on seed complementarity alone may be improved by including information from the flanking regions.

## 4 Conclusion

On model organisms (H. *sapiens, M. musculus, D. melanogaster*) and across validation techniques or localisations of miRNAs/mRNAs, we illustrated the consistency of and specific variations in targeting rules (discriminative features). Similarly as in plants [11], we have shown that flexible sets of rules for predicting interactions may increase the accuracy and sensitivity of classifiers. An essential asset of *fea*miR, derived from the Forward Feature Selection and the embryonic Genetic Algorithm, is the prioritisation of features which may be linked to yet unknown biological, mechanistic details.

To reduce the high false-positive rates, we extended our focus outside the complementary region, to the flanking regions. Across cell types, we analysed the variation in the number of significant features against the distance from the complementary region. We observed a large number of significant features on the upstream and downstream flanking regions, with significance comparable to seed and compensatory regions and a decrease in number and significance levels as we move away from the miRNA complementary region, with a clear drop-off after 40nt across cell types.

## 5 Methods

Using miRNA and 3’UTRs as input, corroborated with information on wet-lab validated interactions or localization, we developed *fea*miR for determining discriminative sets of features that may increase the accuracy/sensitivity of predictions of miRNA/mRNA interactions. These classifiers were tested on various tissues and organisms; both consistent features and subtle differences were described.

On the sets of positive (validated) and negative (non-validated) interactions (generated as described in Supplementary method 1), we first applied statistical tests to determine discriminative features. All feature-frequencies were scaled to 100; *χ^2^* and Fisher exact tests were used to compare the frequency-distributions and to assess the individual contribution of each level of the categorical variables, respectively. For all statistical tests, significance thresholds of 0.1 and 0.2 for marginal significance, under Benjamini-Hochberg corrections, are accepted [26].

We also tested **Machine Learning (ML) approaches**: Decision Trees (DTs), Random Forests (RF) and Support Vector Machines (SVMs), enriched with adapted feature selection approaches. To assess the stability of ML models and evaluate the bias-variance trade-off, we used k-fold *(k*= 10) cross-validation.

First, trained **DTs** were used for feature selection; we maintained a record of frequently-used features and their depth in the DT (restricted to the first 10 levels [27]). Over 100 runs, we retain the feature frequencies, which are subsequently sorted in descending order. The most frequently chosen features are selected (further details in Supplementary method 2).

Secondly, from the **RF** models we retained the variable/feature entropy-based importance assessed on mean decrease in node impurity (Gini indices) and mean decrease in accuracy i.e. the difference in prediction error between each tree and the tree without the focus-feature). The error across trees is averaged and normalised on the standard deviation of the differences (further details in Supplementary method 3).

For the SVMs, we employed Forward Feature Selection (FFS, Supplementary method 4, algorithm 3), a novel version of the classical forward selection used for linear models, augmented with a *nextbestfeature* selection algorithm (Supplementary method 4, algorithm 4). The output is the list of all features, ordered by discriminative power.

### 5.1 Feature selection using GAs

We also propose an optimised approach, based on Genetic Algorithms (GAs, Supplementary method 5), to search for an optimal subset of features through a vast combinatorial set. A set of features is encoded in a binary vector, called a chromosome with a 1 or 0 denoting whether a given feature is included in the feature set. A ‘fitness score’ is defined per chromosomes based on its accuracy. To overcome the limitations of standard GAs, we used an embryonic Genetic Algorithm for feature selection (eGA [22], shown in Supplementary method 6).

Briefly, we maintain an ongoing set of ‘good’ features which is iteratively improved by randomly selecting new features from the pool and applying *forwardfeatureselection* to the ‘good’ and newly selected features. A ‘goodness of feature’ list is maintained at every stage, storing the number of times each feature has been included in the *L* features chosen by *forwardfeatureselection.* Crossover and mutation are applied to the L selected features and if they improve the accuracy then they replace the existing L features. The next set of *k — L* features is randomly selected from the remaining *N — L* features, where *N* is the total number of features created at the start of the algorithm. This revised set of features is processed in the next iteration of the algorithm. This continues until we have k ‘good’ features, or some maximum number of iterations have passed, or there have been k — L unsuccessful attempts to improve the set of features.

### 5.2 Processing of the single cell smartSeq data

All samples were evaluated using MultiQC; for the sRNA samples, the sequencing adapters were trimmed using cutadapt [28]; for the mRNA samples, to balance the distribution of reads across samples/cells, the reads were subsampled without replacement to 3.5M [29]; the sRNA reads were aligned to *H. sapiens* microRNA hairpins and mature sequences [30] using PatMaN, with maximum one mismatch and no gaps (-g 0 -e 1) [31]. The microRNA expression was summarised [32] and the resulting distributions plotted to identify an appropriate signal/noise threshold *(s/n*= 10) [33]. All miRNAs with maximum abundance across samples <10 were discarded.

The preprocessed mRNA samples were aligned to the *H. sapiens* transcriptome (Ensembl 101.38) using STAR [34]. The expression was summarised using feature counts [35]. The signal/noise threshold was determined to be 10 [29]. Next, we evaluated how many of the predicted/validated miRNA/mRNA interactions were supported by miRNAs and mRNAs expressed in the same cell.

## 6 Materials

The miRNA/mRNA interactions for *H. sapiens*, *M. musculus* and *D. melanogaster* were characterised using: miRBase [30] for mature miRNA sequences, Ensembl [36] and FlyBase [37] for reference transcriptomes and miRTarBase [20] for validated interactions, split per validation approach (immunoprecipitation, qRT-PCR, in-situ hybridization and luciferase).

For further assessing predicted miRNA/mRNA interactions, we processed a smartSeq co-sequencing single-cell dataset, GSE114071 [25]; from both assays we retained the presence/absence information for the mRNAs and miRNAs, respectively.

## Acknowledgements

E.W. and I.M. acknowledge the constructive feedback from the Core Bioinformatics group, Cambridge Stem Cell Institute. All authors would also like to acknowledge the use of the University of Oxford Advanced Research Computing (ARC) facility in carrying out this work (http://dx.doi.org/10.5281/zenodo.22558).

## Contributions

E.W., A.C. and I.M. participated in designing the study and analyzing the data. E.W. and I.M. wrote the manuscript. All authors read and approved the submitted manuscript.

